# Validation of an ultra-fast CNV calling tool for Next Generation Sequencing data using MLPA-verified copy number alterations

**DOI:** 10.1101/340505

**Authors:** Bas Tolhuis, Hans Karten

## Abstract

DNA Copy Number Variations (CNVs) are an important source for genetic diversity and pathogenic variants. Next Generation Sequencing (NGS) methods have become increasingly more popular for CNV detection, but its data analysis is a growing bottleneck. Genalice CNV is a novel tool for detection of CNVs. It takes care of turnaround time, scalability and cost issues associated with NGS computational analysis. Here, we validate Genalice CNV with MLPA-verified exon CNVs and genes with normal copy numbers. Genalice CNV detects 61 out of 62 exon CNVs and its false positive rate is less than 1%. It analyzes 96 samples from a targeted NGS assay in less than 45 minutes, including read alignment and CNV detection, using a single node. Furthermore, we describe data quality measures to minimize false discoveries. In conclusion, Genalice CNV is highly sensitive and specific, as well as extremely fast, which will be beneficial for clinical detection of CNVs.

## Introduction

DNA Copy Number Variations (CNVs) are an important source for genetic diversity and significantly contribute to pathogenic variants. It has been demonstrated that CNVs present in the germline DNA contribute to human diseases such as, but not limited to, neurodevelopmental disorders (Wilfert et al 2017), Parkinson’s disease (La Cognata et al 2016), ALzheimer’s disease (Cuccaro et al 2017) and cancer susceptibility (Krepischi et al 2012).

Clearly, accurate and comprehensive detection of CNVs is clinically important. In recent years, Next Generation Sequencing (NGS) methods have become increasingly more popular for CNV detection. Compared to other CNV detection methods, NGS is able to detect variations with higher coverage and resolution, better detection of breakpoints, and better capability to identify novel CNVs (Cuccaro et al 2017). It is thought that NGS will replace other methods in time, when its costs are further reduced and turnaround times of NGS experiments are reduced (Nowakowska 2018).

Computational analysis and storage of NGS data are indispensable for detection of genetic variation and therefore play a role in the total costs and turnaround times of NGS experiments. Over the past few years, we witnessed reducing sequencing costs and increasing amounts of data, which indicates that scalable and cost effective computational solutions are required now and in the near future (Muir et al 2016).

Genalice develops NGS secondary analysis solutions that successfully take care of turnaround time, scalability and cost elements associated with NGS computational analysis, without sacrificing quality. Genalice solutions include read alignment and germline, population and somatic variant calling. Due to its high processing speed, Genalice software has successfully been deployed in a clinical setting to assist in the rapid pediatric sequencing and diagnosis of critically ill infants (Boukhibar et al 2018). Compared to other computational pipelines, Genalice software solution is much faster and requires less storage per data sample, while being highly accurate in detecting Single Nucleotide Variants (SNVs) and insertions/deletions (Pluss et al 2017).

Genalice CNV is a novel tool for fast and accurate detection of CNVs. It uses coverage of two samples, or a sample and a control group of samples to calculate a normalized differential between the two inputs (divided in 100 base buckets), the so-called CNV ratio. In case of a control group, it calculates a standard deviation per bucket of the group to create a statistical background for gain (duplication), loss (deletion) or normal assessment for a segment of the genome.

We assessed the accuracy of Genalice CNV by analyzing a recently published dataset: the ICR96 Exon Validation series (Mahamdalie et al 2017). This dataset consists of high quality sequencing data from a targeted NGS (cancer) assay comprising 96 independent samples. Moreover, the samples have been analyzed with Multiplex Ligation-dependent Probe Amplification (MLPA), resulting in 66 samples with at least one validated CNV (true positives) and 30 samples with validated true negative results in 26 genes. These MLPA-verified results allow for orthogonal evaluation of the calls made by Genalice CNV and a reliable assessment of its accuracy.

We further demonstrate several data quality controls that guard reliable CNV calling. Genalice CNV is highly sensitive with a true positive rate of 98.4%, while calling only a few false positives (n=14 false positive rate of 0.8%). Finally we show that Genalice CNV is extremely fast. It requires less than 15 minutes to call CNVs across all ICR96 samples. Even when alignment of the reads is included, the total processing time is less than 45 minutes for all samples using a single compute node.

## Results

### Accurate CNV calling requires proper data quality control

We observed that several ICR96 samples show an exceptionally high count of buckets with CNV calls (Figure 1A). Up to 70% of the TruSight Cancer Panel captured regions were called as CNV (??). This is an unusual result and we investigated the underlying cause.

**Figure 1:**
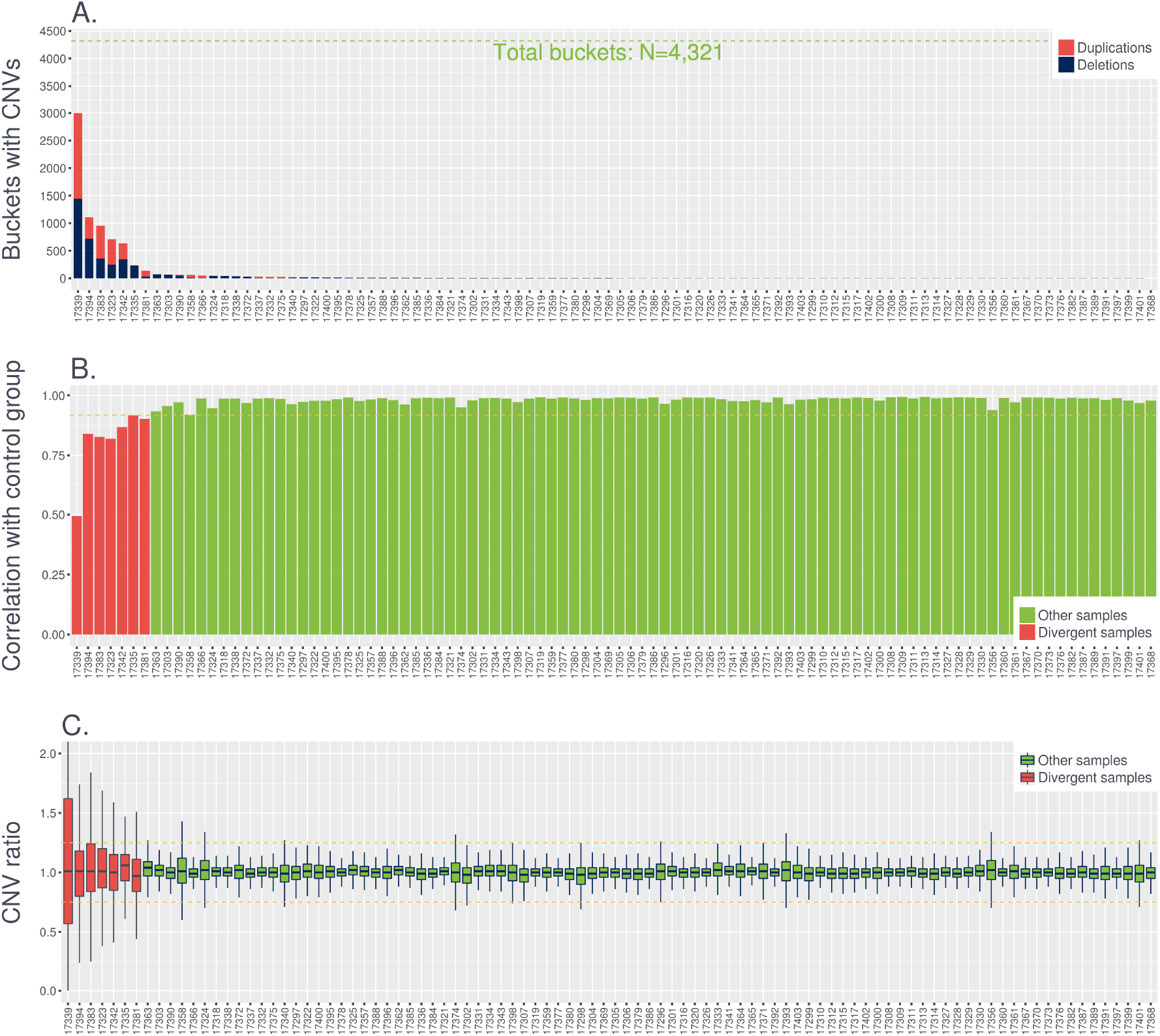
Sample effects cause outlier results. **A**. Number of segments or with a called CNV for all 96 samples. Samples are ranked from highest (left) to lowest number of CNV buckets. CNV calls can be either deletions (blue) or duplications (red). The TruSight Cancer Panel captured regions overlap with 4,321 buckets (green dashed line). **B**. Pairwise correlations of coverage depths between all target 96 samples and their control group. The seven divergent samples, with the highest CNV counts, are indicated (red) and all seven have lower correlation coefficients. The yellow dashed line indicates the sample filter threshold (r=0.9175). Samples with lower coverage correlations are discarded from further analysis. **C**. Box plots contain CNV ratios across all buckets for all 96 samples. Samples are ranked as in A. In addition, the seven divergent samples, with the highest CNV counts, are indicated in red and other samples in green. Yellow dashed lines are CNV significance thresholds.

We first inspected coverage depths and noticed that these seven samples havelower correlation coefficients with their control group (Figure 1B). This leads to CNV ratios and Z scores that extend well beyond the significance thresholds (Figure 1C and data not shown), which explains the high number of CNV calls. Visual inspection showed that CNV calls are irregular in those samples and buckets with duplication, deletion and normal calls alternating within single genes (Figure S1). Taken together, these results suggest that divergent coverage (leading to altered and skewed CNV ratios and Z scores and yielding an unusual high number of CNVs) can be indicative of producing false calls.

Indeed, when we compared the initial Genalice CNV calls to the MLPA results we noticed that the seven divergent samples also have higher False Positive Rates (FPRs) (Figure S2, ??). The seven samples have a median FPR of 9.8%, while the median FPR of the other samples is 0%.

We used the correlation coefficient between target sample and control group coverage as filter mechanism to remove low quality samples (??). All seven divergent samples have a correlation coefficient of less than 0.9175.

The ICR96 dataset includes two separate library enrichment pools (Mahamdallie et al 2017) that have different coverage characteristics. When we examine coverage depths of samples from either pool 1 or pool2, we observe better correlations as compared to samples from both pools (Figure S3A). In addition, coverage depths differ slightly between the two pools (Figure S3B) and samples from Pool 1 have less coverage (Median = 503) compared to pool 2 (Median = 538). Enrichment coverage characteristics can influence CNV ratios and z scores. Indeed, we observed biases in CNV ratio (Figure S3C) and Z score (data not shown) when the control group consists of a mix of samples from both pools. The median CNV ratio and Z scores for a mixed control group are not balanced around 1 and 0, respectively, which is to be expected for these data variables and is the case for pool-specific control groups (Figure S3C). Finally, we observed less CNV calls when the control group is a mix of samples from the two pools (Figure S3D), resulting in reduced sensitivity (data not shown). Here, we avoid reduced sensitivity and biases by using a control group consisting of samples that come from the same library enrichment pool as the target sample.

In summary, for our final analysis with Genalice CNV we excluded seven samples with divergent coverage (??) and used library enrichment pool specific control groups to call CNVs.

**Table 1:** Samples filtered out due to divergent coverage depths. Buckets is the number of buckets with a CNV call. The proportion of the TruSight Cancer Panel has a copy number change. Correlations shows Pearson’s correlation coefficient for target coverage and control group average coverage. The buckets were filtered out. MLPA positive highlights MLPA-verified CNV calls that were removed.

| Sample | Buckets | % Capture | Correlation | % FPR | MLPA negative | MLPA positive |
| --- | --- | --- | --- | --- | --- | --- |
| 17323 | 709 | 16.40 | 0.8189 | NA | 0 | NSD1 Exon 1-23 deletion |
| 17335 | 235 | 5.40 | 0.9169 | 7.9 | 1,119 | BRCA2 Exon 14-16 deletion |
| 17339 | 3,010 | 69.70 | 0.4939 | 66.8 | 1,211 | - |
| 17342 | 634 | 14.70 | 0.8681 | 11.8 | 1,130 | BRCA2 Exon 1-11 deletion |
| 17381 | 136 | 3.10 | 0.9020 | NA | 0 | NSD1 Exon 1-2 deletion |
| 17383 | 963 | 22.30 | 0.8273 | 6.5 | 62 | NSD1 Exon 1-5 deletion |
| 17394 | 1,108 | 25.60 | 0.8405 | 28.6 | 1,207 | BRCA1 Exon 1-2 deletion |

### CNV ratios and Z scores are distinct for MLPA verified loci

Genalice CNV divides the genome into consecutive segments or buckets, which are 100 bases wide in this study. For each bucket it calculates a differential between normalized coverage values from the target sample and the control group. This is the so-called CNV ratio and bucket with a ratio of 1 means that the target has a similar coverage depth or copy number relative to the control group in this genome segment. Furthermore, Genalice CNV calculates a z score for every bucket using the normalized coverage of the target and the average and standard deviation over the control group of samples. This z score allows filtering out CNV ratios with lower statistical significance.

We first examined whether the CNV ratio and Z score measures are sufficient to separate CNV from normal calls. For every sample, we grouped all buckets according to MLPA results into deletion, duplication and normal categories. The resulting density distributions show that CNV ratios and Z scores are distinct for the three categories (Figure 2). For example, buckets that overlay with MLPA normal loci have a median CNV ratio of 1, whereas deletions and duplications have median ratios of 0.59 and 1.36, respectively. We noticed similar differences for Z score values where the median values were 0 (normal), −6.80 (deletion) and 6.46 (duplication).

**Figure 2:**
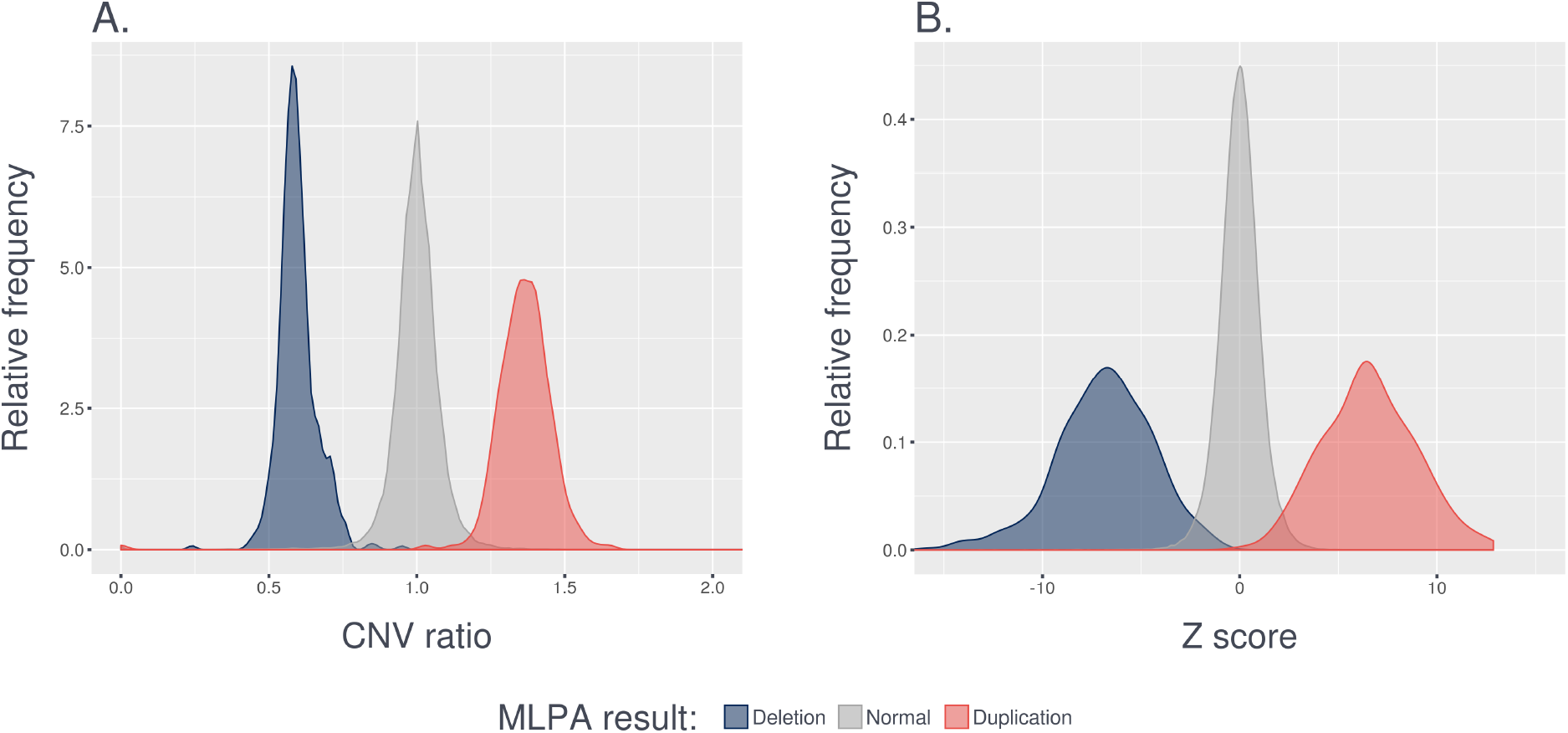
CNV ratios and Z scores are distinct for MLPA verified loci. **A**. Density distribution shows CNV ratios for buckets that overlap with MLPA deletion (blue), MLPA normal (gray) and MLPA duplication (red) results. **B**. Same density distributions as in A, but now for Z scores.

We based our CNV calling thresholds on the observed CNV ratio and Z score distributions. Genalice CNV calls a deletion when a bucket has a CNV ratio less than 0.75 and a Z score less than −4.7, while a duplication requires a CNV ratio higher than 1.25 and a Z score higher than 4.7. Buckets that do not exceed these thresholds are called to have a normal copy number. Buckets that lack coverage in the control group are filtered out. Consecutive buckets with the same call are concatenated into a larger region and can (for this dataset) maximally span the length of an exon.

This Genalice CNV configuration resulted in 415 exons with a CNV call (Table S1), including 275 deleted and 140 duplicated exons. There were 24 samples without CNV calls. Next, we compared the Genalice CNV results to the MLPA results in order to determine the accuracy of our CNV calling tool.

### Genalice CNV is highly sensitive and specific

Using the MLPA results as truth set, we can determine the number of true positive, false negative, true negative and false positive calls in the Genalice CNV NGS results. A true positive (TP) call is defined as a Genalice CNV call that overlaps an MLPA CNV call. Furthermore, both methods agree on the type of CNV being either deletion or duplication. A false negative (FN) is an MLPA CNV call that is not present in the Genalice CNV call set. A true negative (TN) means that both MLPA and Genalice CNV call a normal copy number for all exons that make up a single gene. Finally, false positives (FPs) are Genalice CNV calls that occur in genes that are considered to have a normal copy number based on the MLPA results.

Genalice CNV calling is highly sensitive and specific (Table 2 and Table S2). It correctly calls 62 out 63 MLPA verified CNVs, giving a sensitivity or True Positive Rate (TPR) of 98.4%. In addition, the vast majority of other calls were true negative (1,637 out of 1,651 genes), which equates to a specificity of 99.2% or a False Positive Rate (FPR) of 0.8%. TP and TN Genalice CNV results have clear CNV ratios and Z scores. Where TPs exceed our thresholds and true negative exons have values within the boundaries of our thresholds (Figure 3A-C).

**Figure 3:**
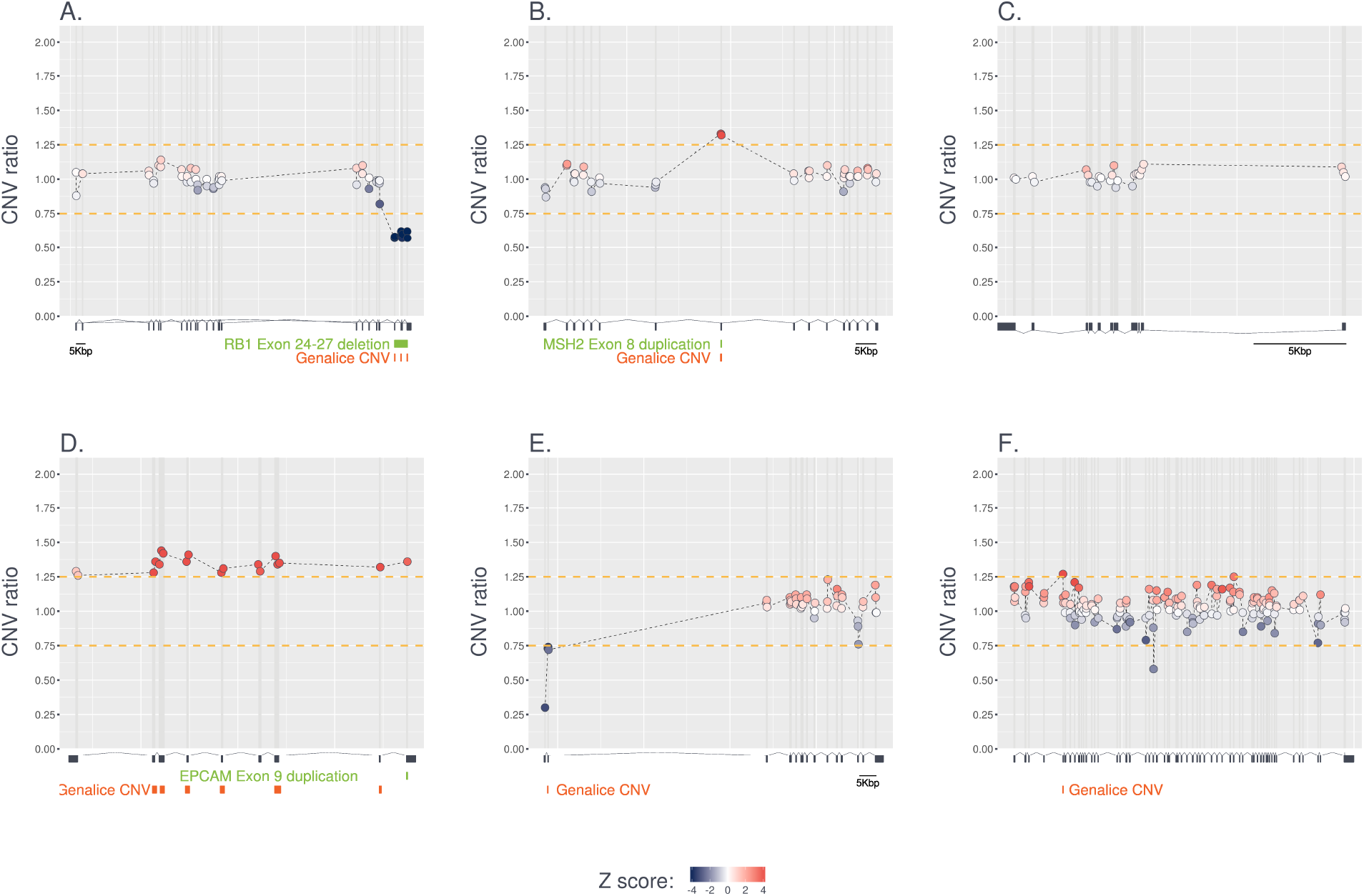
Genalice CNV results for several example genes. **A**. True positive deletion of RB1 exons 24 through 27 in sample 17336. **B**. True positive duplication of MSH2 exon 8 in sample 17316. **C**. True negative TP53 gene in sample 17402. **D**. False negative CNV call in EPCAM gene of sample 17366. MLPA showed that exon 9 is duplicated. This call is not made by Genalice CNV. Instead, the tool calls duplications for exons 2 through 8. The CNV ratio in exon 9 is above the threshold, but the z score is not significant. **E**. False positive deletion at the 5’ end of the CDH1 gene in sample 17303. **F**. False positive duplication in the ATM gene (sample: 17340). **A-F**. For each bucket the CNV ratio is plotted. A ratio of 1 means that the target sample copy number is equal to the average in the control group. Yellow dashed lines show CNV ratio thresholds: 0.75 (deletions) and 1.25 (duplications). Buckets are filtered out when their z score value does not exceed the z score thresholds: −4.7 (deletions) and 4.7 (duplications). Color gradient shows z score value of each bucket. Exon positions are shown below each graph. Connecting lines indicate introns. Connecting lines at the top indicate that the gene is transcribed from the DNA plus strand, while lines at the bottom line show transcription from the minus strand. MLPA-verified CNVs are highlighted in green. Genalice CNV calls are highlighted in orange.

**Table 2:** Genalice CNV accuracy results. Abbreviations in column names stand for: TP = true positives; FN = false negatives; TPR = true positive rate; TN = true negatives; FP = false positives; FPR = false positive rate.

| Data (sub)set | TP | FN | % TPR | TN | FP | % FPR |
| --- | --- | --- | --- | --- | --- | --- |
| Pool 1 | 31 | 0 | 100.0 | 827 | 11 | 1.3 |
| Pool 2 | 31 | 1 | 96.9 | 810 | 3 | 0.4 |
| ICR96 | 62 | 1 | 98.4 | 1,637 | 14 | 0.8 |

The FN call is a missed exon 9 duplication of the EPCAM gene in sample 17366 (Figure 3D). However, Genalice CNV does detect duplications in four other exons and CNV ratios and z scores are elevated throughout the EPCAM gene in this sample. As such, the duplication in the EPCAM gene will not go unnoticed when using Genalice CNV for detecting copy number variations. The CNV ratio in exon 9 is 1.36 and well above our threshold. Yet, exon 9 is filtered out, because its z score is 3.74 and below our selected significance threshold of 4.7.

Figure 3E-F show examples of FP calls. Most FP CNV calls (11 out of 14) are detected in Pool 1 (Table 2), which has lower coverage depth on average (Figure S3). This is a striking difference, because the number of TN genes is more or less equal between the two pools. Nine FP genes have a CNV ratio between 0.7 and 1.3 (i.e. close to our thresholds of 0.75 and 1.25). Their z scores range between −4.9 and −6.1 for deletions and 4.9 and 6.1 for duplications. Only 3 TP have ratios in this window. Clearly, the majority of FP calls consist of weaker evidence compared to TP calls. Genalice CNV calls 4 FP exons in sample 17303. Interestingly, all 4 exons have a GC content higher than 60%. The FPs include eleven different genes: APC, ATM (2x), BARD1 (2x), BRCA2, BRIP1, CDH1 CDKN2A, PTEN (2x), RAD51D, SMAD4 and STK11.

### Genalice Map and CNV are extremely fast

In addition to highly accurate CNV calls, Genalice software provides extremely fast read alignment and CNV calling of the ICR96 dataset (Table 3). Genalice Map aligned reads in 17 seconds per sample on average and the entire cohort took less than half an hour. Genalice CNV requires a control group and to this end we created a Genalice Coverage (GCO) files for each library enrichment pool. A GCO file contains averages and standard deviations of the group for every bucket in the genome, which can be used to calculate z scores. Making a GCO file for a single pool (N=45 samples) took 81 seconds for pool 1 and 78 seconds for pool 2. This equates to an average of less than 2 seconds per sample. Finally, Genalice CNV required approximately 7 seconds per sample to call CNVs.

Genalice software processed the entire ICR96 dataset in less than 45 minutes from input sequence reads (FASTQ) to high quality CNV calls.

**Table 3:** Genalice runtimes for processing the ICR96 dataset. Time is displayed in HH:MM:SS. Genalice Map indicates turnaround times of read alignment. Genalice CNV (control group) gives times required to create coverage depth averages and standard deviations of a set of control samples. Genalice CNV (call CNVs) reports the time required to make CNV calls.

| Process | Average (N=1) | Sum (N=96) |
| --- | --- | --- |
| Genalice Map | 00:00:17 | 00:27:08 |
| Genalice CNV (control group) | 00:00:02 | 00:02:39 |
| Genalice CNV (call CNVs) | 00:00:07 | 00:11:55 |
| Complete workflow | 00:00:26 | 00:41:41 |

## Discussion

Genalice CNV is a highly accurate copy number detection tool, which operates at extremely high speed. To warrant high accuracy, we had to take into account quality aspects of the NGS data, including: good correlation between the target sample coverage and control group average coverage; and library enrichment pool specific control groups. If a target sample does not adhere to specific quality constraints, then false positive rates might increase.

The library enrichment pool specific control group seems a limitation for using NGS data for CNV detection, because it will be challenging to pick a fixed set of samples as a control group that can be used independent of enrichment batches. However, it remains to be seen whether this limitation is either a general feature of targeted NGS assays or specific for the TruSight Cancer Panel assay used in this study. It is known that various capture technologies differ in enrichment efficacy, coverage in specific genome regions, and coverage uniformity (Chilamakuri et al 2014, Asan et al 2011)

Currently, Genalice CNV quantifies copy number changes using read depth information. Such approach fits well with targeted NGS assays, because the break points flanking the copy number change are (unless part of the panel design) likely to occur outside the captured DNA sequences. This makes split-read and paired-end detection methods less suitable for targeted assays. However, NGS CNV detection based on read depth might suffer from bias due to GC content (Xi et al 2016, Talevich et al 2016) and capture kit specific output variation. We observed FP CNVs in sample 17303 in exons with high GC content. Nevertheless, filtering based on GC content will not be trivial, because more than 10% of the TN exons have a similar high GC content.

With an emerging use of NGS in clinical diagnostics scalability and data processing speed become more important. Genalice processes the entire ICR96 Exon Validation series of 96 targeted NGS assays within 45 minutes. This includes read alignment, creating control groups and calling CNVs across all samples. CNV calling alone takes less than 15 minutes for 90 samples. To the best of our knowledge, the DECoN tool is the only other tool that has been validated using the ICR96 Exon CNV validation series. DECoN called CNVs across 48 samples of the same dataset in 24 minutes (Fowler et al 2016). We don’t know the actual hardware configuration that DECoN used, which makes a direct comparison with Genalice CNV somewhat challenging. However, it is clear that both tools are sufficiently fast for clinical use. It is important to realize that, the DECoN turnaround time excludes read alignment. We measured Genalice Map to be approximately 40 times faster than the BWA-MEM read aligner. Such processing acceleration can be essential when clinical sequencing is performed at large scale.

We used the orthogonally verified CNVs (Mahamdalie et al 2017) to obtain an unbiased and reliable accuracy measure for Genalice CNV. The DECoN tool was validated using the same ICR96 Exon Validation series, but results of only 48 samples are presented (Fowler et al 2016). This tool detected 16 out of 16 BRCA exon CNVs. Likewise, Genalice CNV correctly called 22 out of 22 BRCA exon CNVs. Amongst the other 15 genes, DECoN missed an exon 22 deletion in NSD1. Similarly, Genalice CNV missed an exon 9 duplication in EPCAM. These results demonstrate that Genalice CNV and DECoN have similar CNV calling sensitivity with the ICR96 Exon Validation series.

Specificity is equally high and around 99% for both tools. For example, DECoN detected one BRCA CNV that was not confirmed by MLPA (Fowler et al 2016). Genalice CNV also reported one false positive BRCA exon CNV.

The ICR96 Exon Validation series has sequencing data generated using the TruSight Cancer Panel (Mahamdalie et al 2017) and its average coverage depth per exon per sample is relatively high (500x). Typical whole exome sequencing (WES) and whole genome sequencing (WGS) assays have lower coverage depths. This raises the question how accurate Genalice CNV is with lower coverage depths. Of course, this requires additional validations of the tool. However, for the ICR96 true negative and true positive exons, coverage depth ranges from 0 to more than 1700x, indicating that Genalice CNV correctly calls copy numbers over a wide range of coverage depths including lower coverage depths.

The ICR96 Exon Validation series allowed us to benchmark the Genalice CNV tool in an independent and reliable manner. It is demonstrated that Genalice CNV is fast and accurate when detecting CNVs in a targeted NGS assay. Unlike many other CNV tools, Genalice CNV is applicable to multiple NGS assays, including targeted, whole exome and whole genome sequencing methods.

## Materials and methods

A full description of the material and methods can be found on our github page: https://github.com/Genalice/genalice-icr96.

### ICR96 Exon CNV Valadation series

The ICR96 Exon CNV Validation series can be accessed through the European-Genome phenome Archive (EGA) under the accession number EGAS00001002428. Details of how to access the data is available at EGA or from the ICR website: https://www.icr.ac.uk/icr96.

### Reference sequence and alt file

We aligned reads and called CNVs against the GRCh37 assembly of the human reference genome.

This reference assembly contain two types of contigs, namely: primary chromosomes and alternate loci. Genalice Map is an alternate loci aware read alignment tool. Discrimination between primary chromosomes and alternate loci can be achieved through so called alt files. An alt file is included as annotation, while making a Genalice Reference File (GRF). The alt files for the GRCh37 and GRCh38 assemblies can be found on our github page.

### Genalice version

Throughout this study we used Genalice version 2.6.

### Configuration files

Genalice tools are fully configurable to obtain the best possible data results. As a consequence, each tool has a series of options to allow customized settings. Adding multiple options to the command line can be cumbersome. Therefore, multiple option settings can be added as a single configuration file through the `cmd_file` option.

The configuration file(s) used in this study can be found on our github page.

### Open source tools

In addition to Genalice solutions, we used the following open source tools:

- bedtools v2.20.1
- R version 3.3.2

### Hardware configuration

Throughout this study we used a server with the following hardware configurations: a dual Intel Xeon E5 2620 V2 (2100-2600 Mhz) CPU with a total of 12 cores and 24 threads, 128GB RAM (8×16GB, 1333 Mhz), InfiniBand adapter of 40 Gigabit/second to stream data from storage disks to compute nodes. The data is stored on Raid 50 disks with 1GB/second IO speed (read/write). Operating system is SuSE Linux Enterprise Server 12 on standard server board.

### Genalice reference file

Genalice Map requires a reference index for read alignment. To this end, we first made a Genalice Reference File (GRF) from fasta sequences. This GRF is a binary reference sequences, which is essential to the Genalice Aligned Reads (GAR) file format. We annotated the GRF into primary assembly and alternative contigs using a so called alt file. Next, we made an comprehensive Genalice Index File (GRI) using GRF as input.

The exact commands and configurations used in this study can be found on our github page.

### Genalice Map

Genalice Map aligns short sequence reads to the reference index (GRI) with a patented high performance mapping algorithm (REF). First, the GRI is loaded into memory (RAM) and subsequently raw sequence reads of a single sample are matched to the index in memory. After all sequence reads have been processed the alignment result is written to disk in the GAR file format. This mapping procedure repeats itself for every sample of the ICR96 dataset, while the GRI remains in memory.

The read alignment tool is set up for optimal results using the configuration files described above. The exact commands and configurations used in this study can be found on our github page.

### Genalice CNV

In this study, Genalice CNV detects copy number changes in a target sample relative to a control group. The tool divides the genome into consecutive buckets and copy numbers are assessed for each bucket. Here, we used buckets with a length of 100 bases, which is the maximum resolution of the tool. The tool can assign a normal, loss (deletion) or gain (duplication) state to each bucket. Adjacent buckets with identical states are concatenated into larger events. Buckets without sufficient coverage in the control group are filtered out (drop) and no call is made.

#### Control group

The control group is summarized in a Genalice Coverage (GCO) file, containing average coverage depths and standard deviations for every bucket. The average values are used to calculate the CNV ratio and, together with the standard deviation, z scores to assess statistical significance. In order to include samples with varying coverage depths, each sample is first normalized to an arbitrary coverage depth. In this study, the normalised depth is set at 75x.

A GCO file requires GAR files from the control samples as input. In this study, the control samples differ per target sample. First, control and target samples all come from the same library enrichment pool. The ICR96 Exon CNV Validation series is somewhat unusual in its composition, because it has a relatively high number of specific CNVs in a small number of samples. Such composition is highly unlikely in clinical testing (Fowler et al 2016). Furthermore, it may skew averages and standard deviations in the GCO file. So, secondly, we attempted to select MLPA-normal controls for every sample with an MLPA-positive CNV. Finally, we excluded the target sample from the control group. A full list of control group samples can be found on our github page.

#### CNV calling

Genalice CNV requires a target sample (GAR format) and control group (GCO format) to call copy number alterations in the target relative to the control group. It first calculates coverage ratios for the target sample and control groups for every bucket, separately. Coverage ratios are the bucket coverage divided by the sample/control group average coverage. This normalization procedure allows direct comparison of target and control group. Next, the tool calculates a CNV ratio for every bucket by dividing the target coverage ratio over the control group ratio. The tool calls duplications when the CNV ratio was equal to or higher than 1.25. Likewise, it calls deletions whenever the CNV ratio was equal to or less than 0.75. Any buckets with CNV ratios in between those thresholds are considered to have normal copy numbers. Moreover, the tool calculates a z score for every bucket, which can be used to filter out deletion or duplication calls that deviate to little from the control group. Here, buckets with z scores between −4.7 and 4.7 are considered to have normal copy numbers.

The exact commands and configurations used in this study can be found on our github page.

## Acknowledgements

This study makes use of the ICR96 exon CNV validation series data generated by Professor Nazneen Rahman’s team at The Institute of Cancer Research, London as part of the TGMI.

## Supplementary figures

**Figure 4: S1.**
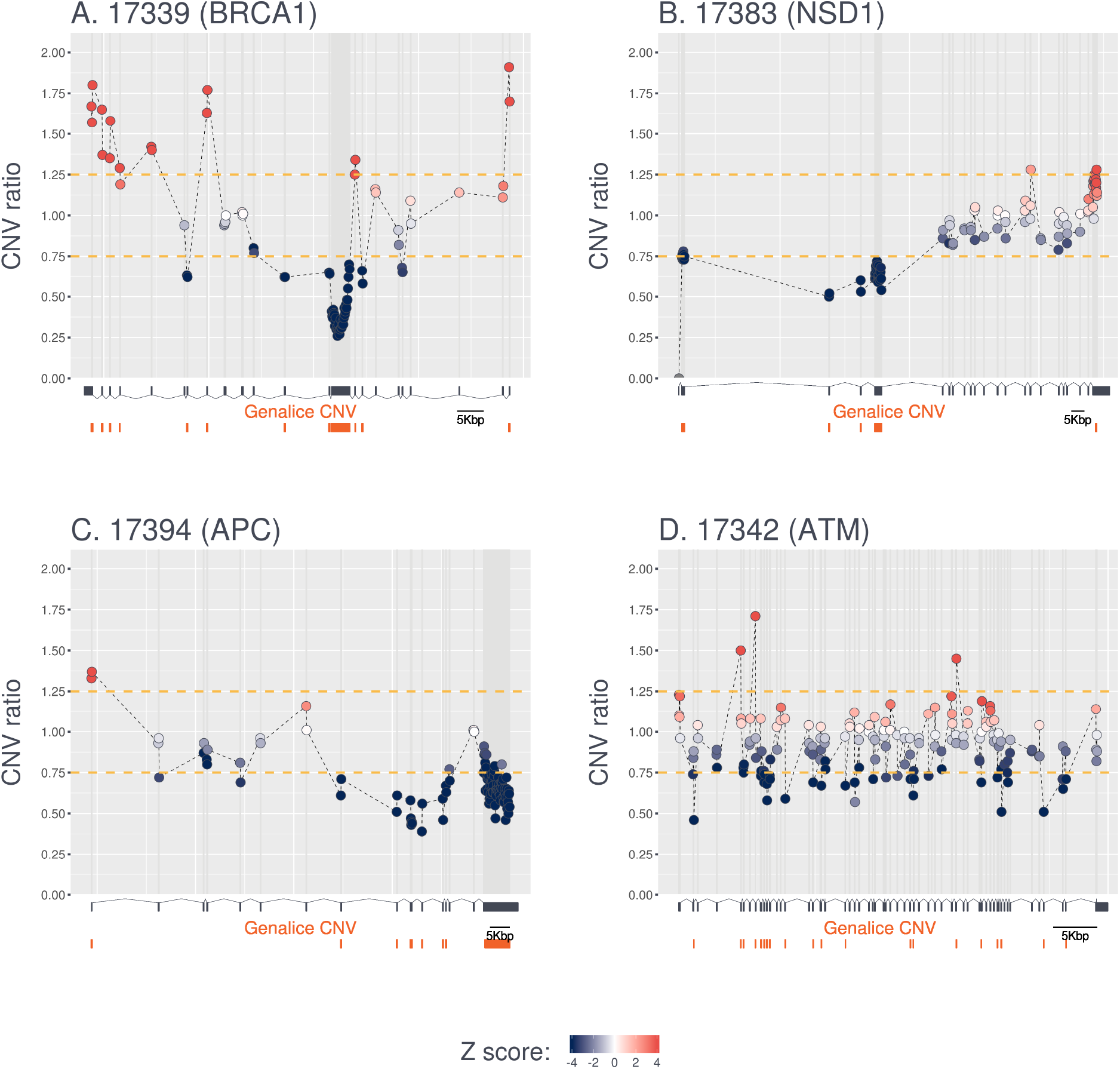
Examples of CNV ratios across genes in four samples with divergent coverage depth. Sample and gene names are shown above each graph. Legend is as described in Figure 3.

**Figure 5: S2.**
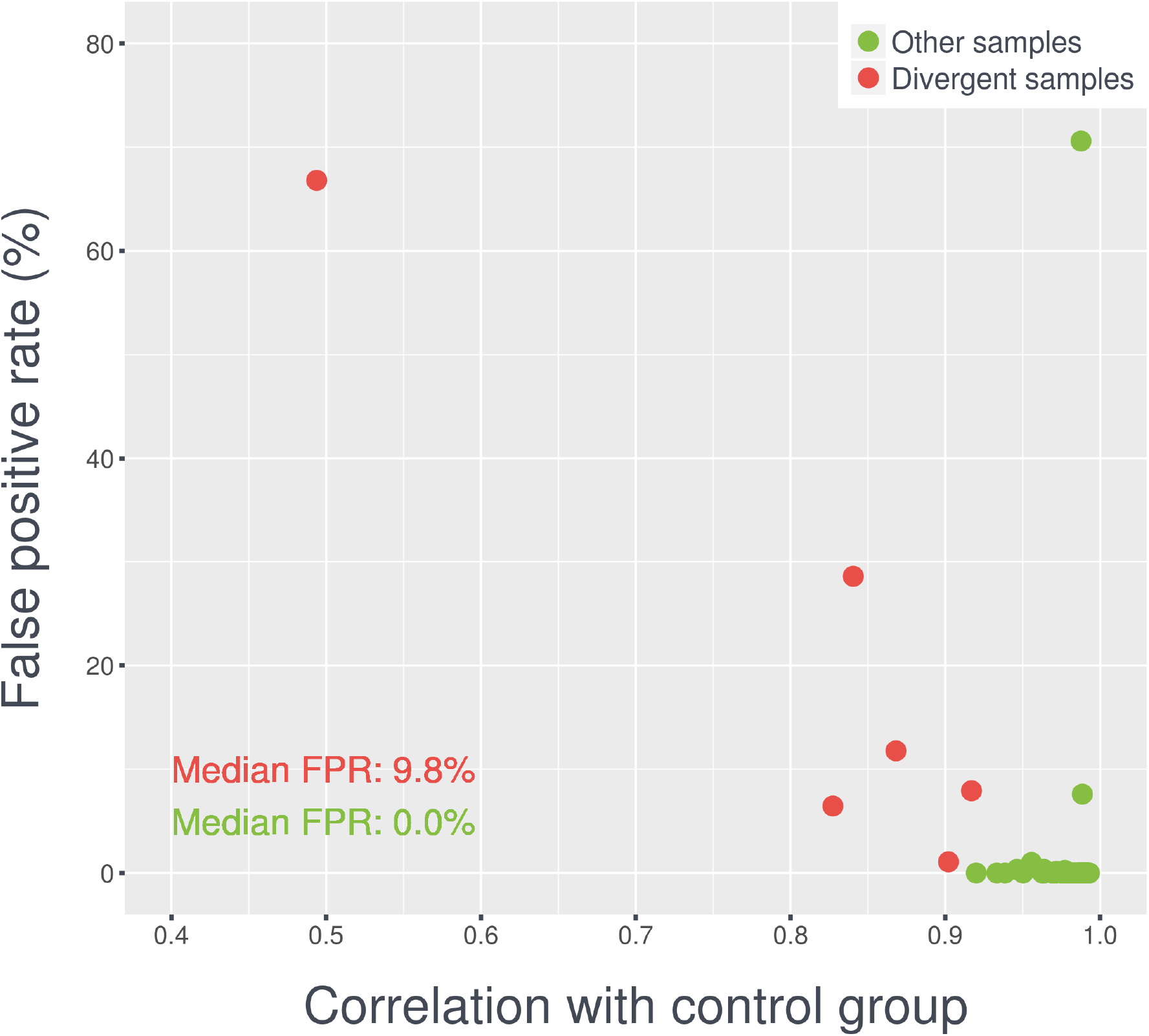
Correlations bewteen target and control group coverage values in relationship to false discovery rates.

**Figure 6: S3.**
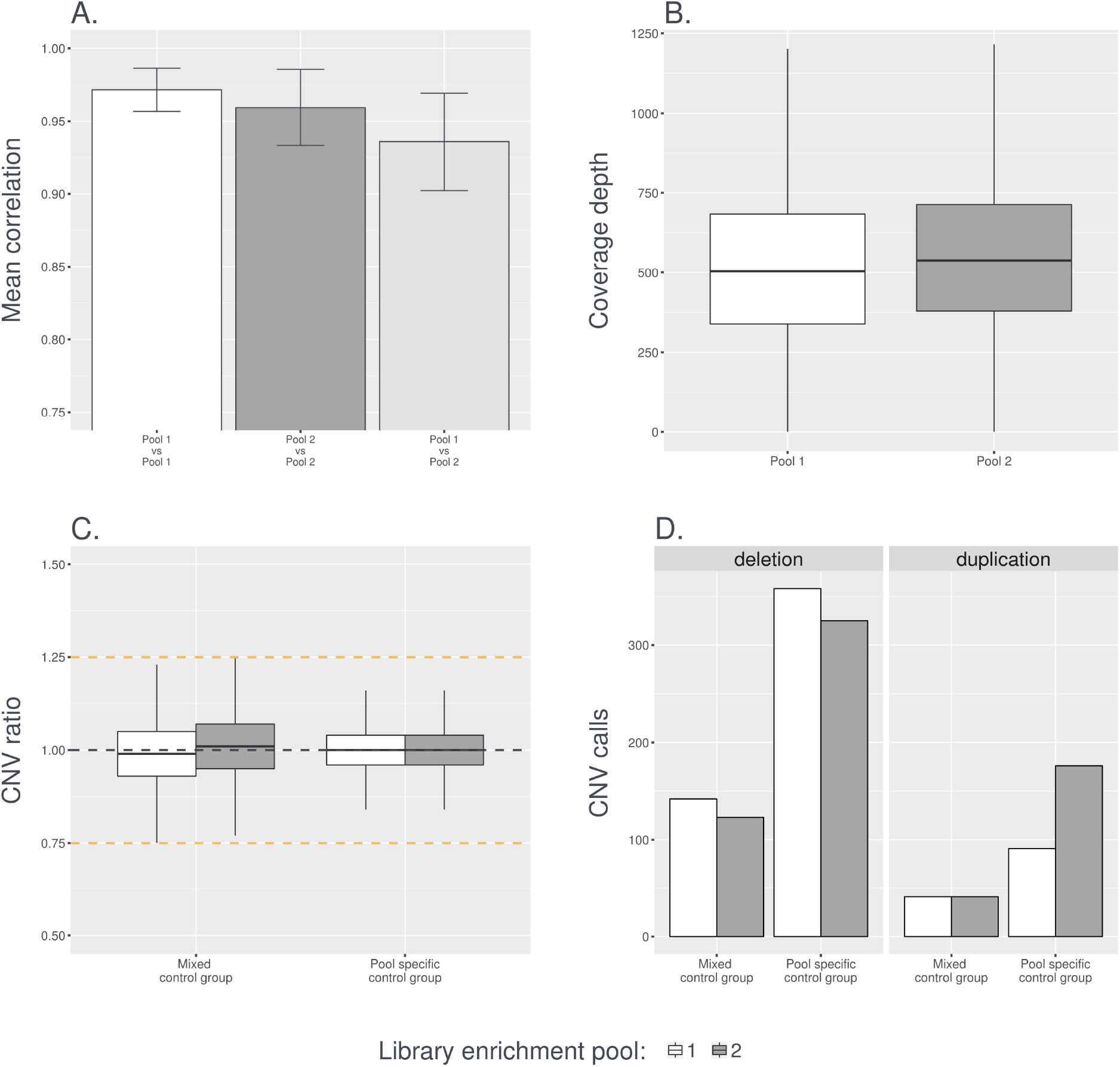
Library enrichment pool effects on CNV calling. **A**. Mean correlations is the average correlation coefficient for all pairwise sample combinations. We compared samples from pool 1 against each other, samples from pool 2 against each other or samples from pool 1 against pool 2. **B**. Boxplots show coverage depths accross all buckets and samples in either pool 1 or pool 2. **C**. Boxplots show distribution of CNV ratios across all buckets and samples in either pool 1 (white) or pool 2 (gray). When a control group with samples from pool 1 and pool 2 is used then the median CNV ratio is at 1. In contrast, when a pool specific control group is used the median is 1. **D**. Barplots show CNV calling yield for deletions (left) and duplications (right). Results are separated in pool 1 (white) and pool 2 (gray). When a pool specific control group is used the yield is much higher.

